# Mutations in *EBF3* disturb transcriptional profiles and underlie a novel syndrome of intellectual disability, ataxia and facial dysmorphism

**DOI:** 10.1101/067454

**Authors:** Frederike Leonie Harms, Katta Mohan Girisha, Andrew A. Hardigan, Fanny Kortüm, Anju Shukla, Malik Alawi, Ashwin Dalal, Lauren Brady, Mark Tarnopolsky, Lynne M. Bird, Sophia Ceulemans, Martina Bebin, Kevin M. Bowling, Susan M. Hiatt, Edward J. Lose, Michelle Primiano, Wendy K. Chung, Jane Juusola, Zeynep C. Akdemir, Matthew Bainbridge, Wu-Lin Charng, Margaret Drummond-Borg, Mohammad K. Eldomery, Ayman W. El-Hattab, Mohammed A.M. Saleh, Stéphane Beziéau, Benjamin Cogné, Bertrand Isidor, Sébastien Küry, James R. Lupski, Richard M. Myers, Gregory M. Cooper, Kerstin Kutsche

**Affiliations:** Institute of Human Genetics, University Medical Center Hamburg-Eppendorf, Hamburg, Germany; Department of Medical Genetics, Kasturba Medical College, Manipal University, Manipal, India; HudsonAlpha Institute for Biotechnology, Huntsville, AL USA; Department of Genetics, University of Alabama at Birmingham, AL USA; University Medical Center Hamburg-Eppendorf, Bioinformatics Service Facility, Hamburg, Germany; Center for Bioinformatics, University of Hamburg, Hamburg, Germany; Heinrich-Pette-Institute, Leibniz-Institute for Experimental Virology, Virus Genomics, Hamburg, Germany; Diagnostics Division, Centre for DNA Fingerprinting and Diagnostics, Hyderabad, Telangana, India; Department of Pediatrics, McMaster University Medical Center, Hamilton, Ontario, L8N 3Z5, Canada; Department of Pediatrics, University of California, San Diego, CA 92123, USA; Division of Genetics/Dysmorphology, Rady Children’s Hospital San Diego, San Diego, CA 92123, USA; Department of Neurology, University of Alabama at Birmingham, Birmingham, AL USA; Department of Genetics, University of Alabama at Birmingham, Birmingham, AL USA; Department of Pediatrics and Medicine, Columbia University, New York NY 10032, USA; GeneDx, Gaithersburg, MD 20877, USA; Department of Molecular and Human Genetics, Baylor College of Medicine, Houston, TX 77030, USA; Human Genome Sequencing Center, Baylor College of Medicine, Houston, Texas, USA; Cook Children’s Genetic Clinic, Fort Worth, Texas, USA; Division of Clinical Genetics and Metabolic Disorders, Department of Pediatrics, Tawam Hospital, Al-Ain, United Arab Emirates; Section of Medical Genetics, Children’s Hospital, King Fahad Medical City, Riyadh, Saudi Arabia; CHU Nantes, Service de Génétique Médicale, Nantes CEDEX 1, France; INSERM, UMR-S 957, Nantes, France

## Abstract

From a GeneMatcher-enabled international collaboration, we identified ten individuals with intellectual disability, speech delay, ataxia and facial dysmorphism and a mutation in *EBF3*, encoding a transcription factor required for neuronal differentiation. Structural assessments, transactivation assays, *in situ* fractionation, RNA-seq and ChIP-seq experiments collectively show that the mutations are deleterious and impair EBF3 transcriptional regulation. These findings demonstrate that EBF3-mediated dysregulation of gene expression has profound effects on neuronal development in humans.

---

Intellectual disability (ID) is a common phenotype with extreme clinical and genetic heterogeneity. Widespread application of whole-genome and whole-exome sequencing (WES) has tremendously increased the elucidation of the genetic causes of non-syndromic and syndromic forms of ID^1,2^. WES together with the freely accessible tool GeneMatcher (http://genematcher.org) which brings together clinicians and researchers with an interest in the same gene, significantly aid in identifying new disease genes^3^.

We investigated a family with three healthy and two affected children, who both presented with global developmental delay, febrile seizures, and gait instability with frequent falls. WES was performed in both probands and one healthy sibling. We initially hypothesized a Mendelian recessive trait; however, we did not identify any rare, potentially pathogenic biallelic variants in the affected siblings (data not shown). WES data were analyzed for heterozygous variants absent in dbSNP138, 1000 Genomes Project, Exome Variant Server, and ExAC Browser, shared by both affected subjects and absent in the healthy sibling. This analysis identified 16 variants (**Supplementary Table 1**). We used objective metrics from ExAC to prioritize genes intolerant to functional variation (pLI ≥0.9 and high values for Z score) (**Supplementary Table 1**) (http://biorxiv.org/content/early/2015/10/30/030338); five genes were identified with strong selection against various classes of variants for segregation analysis in the family (**Supplementary Table 1**). Four variants were inherited from a healthy parent and/or were present in two healthy siblings (**Supplementary Table 2**). The missense variant c.625C>T [p.(Arg209Trp)] in *EBF3* was confirmed in both affected siblings and was absent in the father and all healthy siblings (**Supplementary Fig. 1a** and **Supplementary Table 2**). In leukocyte-derived DNA from the mother, the Sanger sequence profile showed a very low signal for the mutated base (thymine) superimposed on the wild-type sequence (cytosine) suggesting that she had somatic mosaicism for the *EBF3* variant (**Supplementary Fig. 1a**)^4^. By cloning the mutation-bearing *EBF3* amplicon followed by sequencing of colony PCR products, we confirmed the mother to be a mosaic carrier (18% and 4% of leukocytes and buccal cells, respectively, were heterozygous for the *EBF3* variant) (**Supplementary Fig. 1b**); parenthetically, maternal mosaics are at greater recurrence risk^4^. Given that *EBF3* is intolerant to functional genetic variation (**Supplementary Table 1**)^5, 6^ and the variant p.(Arg209Trp) was computationally predicted to be deleterious (**Supplementary Table 3**), we next submitted *EBF3* to GeneMatcher and were matched with eight other research groups.

In addition to the above family, eight unrelated affected individuals with variants in *EBF3* were identified through WES by groups that independently submitted to GeneMatcher. In addition to c.625C>T [p.(Arg209Trp)], we found the four non-synonymous variants c.196A>G [p.(Asn66Asp)] in subject 4, c.512G>A [p.(Gly171Asp)] in subject 8, c.530C>T [p.(Pro177Leu)] in subject 6, and c.422A>G [p.(Tyr141Cys)] in subject 7, the 9-bp duplication c.469_477dup [p.(His157_Ile159dup)] in subject 10, the nonsense variants c.913C>T [p.(Gln305*)] in subject 3 and c.907C>T [p.(Arg303*)] in subject 9, as well as the splice site mutation c.1101+1G>T in subject 5 (Fig. 1a,b and **Supplementary Table 3**). All variants were predicted to impact protein function (**Supplementary Table 3**), and were absent in 1000 Genomes Browser, Exome Variant Server, and ExAC Browser. All eight additional variants were confirmed to have arisen *de novo* (**Supplementary Table 4**). The *inframe* duplication and the five amino acid substitutions affect residues conserved through evolution and invariant among paralogs (**Supplementary Fig. 2**).

**Figure 1.**
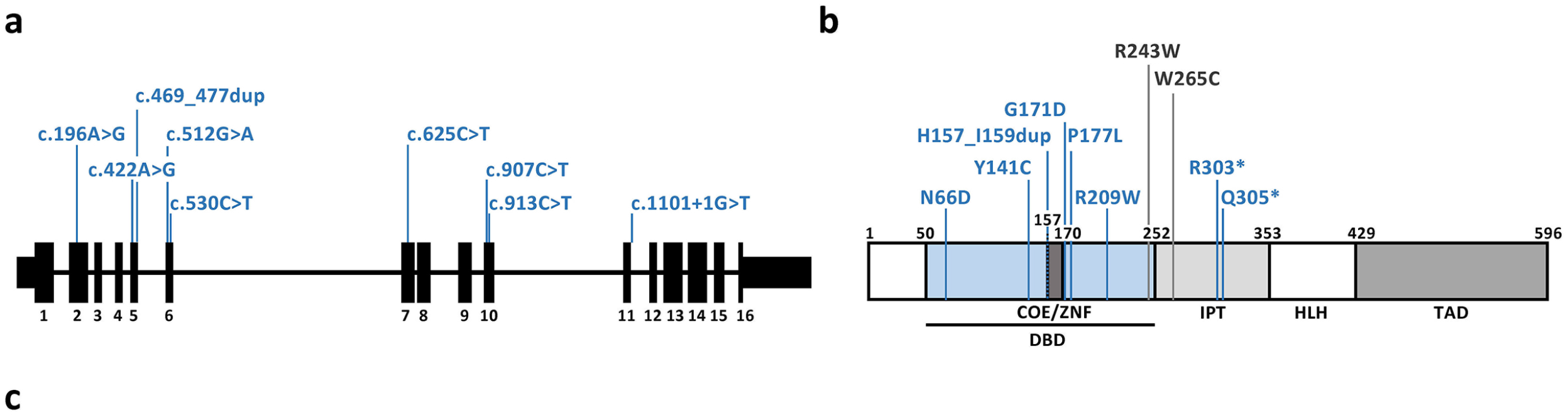
*EBF3* mutations identified in ten patients. (**a**) Schematic representation of the exon-intron structure of *EBF3*. Black bars represent exons and black lines introns. Germline mutations identified in the patients are shown in blue. (**b**) Domain structure of the EBF3 protein with the positions of the identified mutations. *EBF3* germline mutations are indicated in blue and tumor-associated mutations in dark grey. Amino acid numbers are given. DBD: DNA-binding domain with an atypical zinc finger (ZNF; COE motif); IPT: Ig-like/plexins/transcription factors; HLH: helix-loop-helix motif; TAD: transactivation domain. (**c**) Photographs of six patients show subtle, yet distinct facial dysmorphism. Long face, tall forehead, high nasal bridge, deep philtrum, straight eyebrows, strabismus, short and broad chin and mildly dysmorphic ears can be noted in all. Consent for the publication of photographs was obtained for the six subjects.

Consistent clinical features in all individuals with *EBF3* mutation were intellectual disability, speech delay and motor developmental delay (10/10). Ataxia was reported in 6/8 and seizures in 2/9. Brain imaging revealed cerebellar vermian hypoplasia in 2/8 (**Supplementary Table 4**). Facial dysmorphism was mild and commonly seen features include long face, tall forehead, high nasal bridge, deep philtrum, straight eyebrows, strabismus, and short and broad chin (Fig. 1c).

*EBF3* encodes the early B-cell factor 3, which is one of four members of the EBF transcription factor family (also known as Olf, COE, or O/E). All EBFs consist of an N-terminal DNA-binding domain (DBD), an IPT (Ig-like/plexins/transcription factors) domain with yet unknown function, a helix-loop-helix (HLH) domain, which is critical for homo- and heterodimer formation, and a C-terminal transactivation domain (TAD) (Fig. 1b)^7^. EBF1 has been discovered as a key factor for B-cell differentiation^8^ and olfactory nerve signaling^9^. However, expression of *ebf1, ebf2* and *ebf3* in early post-mitotic neurons during embryogenesis suggested a role in neuronal differentiation and maturation^10^. Ebf3 acts downstream of the proneural transcription factor neuroD in late neural differentiation in *Xenopus*^11^ and is transcriptionally repressed by ARX^12^, abnormalities of which cause a spectrum of developmental disorders ranging from ID to brain malformation syndromes^13^. Silencing, genomic deletion and somatic point mutations [p.(Arg243Trp) and p.(Trp265Cys)]^14, 15^ of *EBF3* in diverse types of cancer as well as EBF3-mediated induction of cell cycle arrest and apoptosis suggest that EBF3 acts as a tumor suppressor by regulating the expression of specific target genes and controlling a potential anti-neoplastic pathway^7^. All germline *EBF3* missense variants and the tumor-specific mutation p.(Arg243Trp) are located in the DBD (Fig. 1b and **Supplementary Fig. 2**). Interaction of the DBD with DNA is dependent on a zinc-coordination motif, the zinc knuckle (COE motif), located between His157 and Cys170 (**Supplementary Fig. 2**). Thus, the p.(His157_Ile159dup) variant most likely affects DNA-binding as has been shown for the EBF3^H157A^ mutant^16, 17^.

The structural impact of the five missense mutations was explored by a homology model for the DNA-bound configuration of the DBD of EBF3 with the DNA duplex containing the EBF1 consensus sequence^18^. Two of the five alterations, p.(Asn66Asp) and p.(Gly171Asp), were predicted to directly affect DNA binding (**Supplementary Fig. 3a-c**). Pro177 is localized in close proximity to the zinc finger (His157, Cys161, Cys164 and Cys170) and to Asn174, which forms a hydrogen bond with DNA. Replacement of Pro177 by leucine causes a conformational change, probably affecting correct positioning of the zinc knuckle and destabilizing the protein-DNA complex (**Supplementary Fig. 3d**). Arg209 does not directly interact with DNA, but forms hydrogen bonds with the backbone of Cys198 and Asn197, the latter directly interacting with DNA. The change of Arg209Trp is expected to alter positioning of Asn197 and binding affinity for DNA (**Supplementary Fig. 3e**). Tyr141 is localized within a loop probably involved in EBF3 dimer formation. Substitution of this residue may lead to a conformational change at the dimer interface, resulting in reduced stability of the EBF3 dimer and interfering with its ability to interact with DNA (**Supplementary Fig. 3f**).

To analyze the functional effect of the *EBF3* mutations, all mutant proteins were transiently and efficiently expressed in HEK 293T cells. By confocal microscopy analysis, we confirmed exclusive nuclear localization of wild-type EBF3 (Fig. 2a). In contrast, all but one mutant protein showed nuclear localization or both nuclear and cytoplasmic distribution. Only the mutant lacking 202 amino acids at the C-terminus was almost entirely mislocalized to the cytoplasm (Fig. 2a). We next performed a reporter gene assay to assess whether EBF3 mutants still mediate transcriptional activation of the target gene *CDKN1A* (p21)^16^. Similar to p53, for which *CDKN1A* is a prototypical target gene, expression of wild-type EBF3 markedly increased the reporter activity driven by the *CDKN1A* promoter (Fig. 2b). In contrast, EBF3 mutants had significantly reduced or no ability to activate transcription of the reporter gene (Fig. 2b). *CDKN1A* promoter activation was not significantly reduced when EBF3^WT^ was co-expressed with either of the mutants p.Gly171Asp, p.Pro177Leu and p.Arg209Trp (**Supplementary Fig. 4a**). In contrast, co-expression of wild-type EBF3 and each of the mutants p.Asn66Asp, p.Tyr141Cys, p.His157_Ile159dup, p.Arg303*, and p.Gln305* caused a significant reduction in reporter activity by 40–50% (**Supplementary Fig. 4a**), suggesting a potential dominant-negative impact of these mutations on the wild-type allele. The dominant-negative effect was also observed when expressing increasing amounts of the EBF3 mutant p.Asn66Asp or p.Gln305* in the presence of a fixed amount of wild-type EBF3, demonstrating dose-dependent reduction in reporter gene activity (**Supplementary Fig. 4b**). However, the observed nonsense variants are predicted to undergo nonsense-mediated mRNA decay *in vivo*, and our data also suggest that the truncated protein p.Gln305*, if it was produced *in vivo*, would not localize to the nucleus. Thus, it remains unclear to what extent pathogenesis results from dominant-negative or loss-of-function mechanisms.

**Figure 2.**
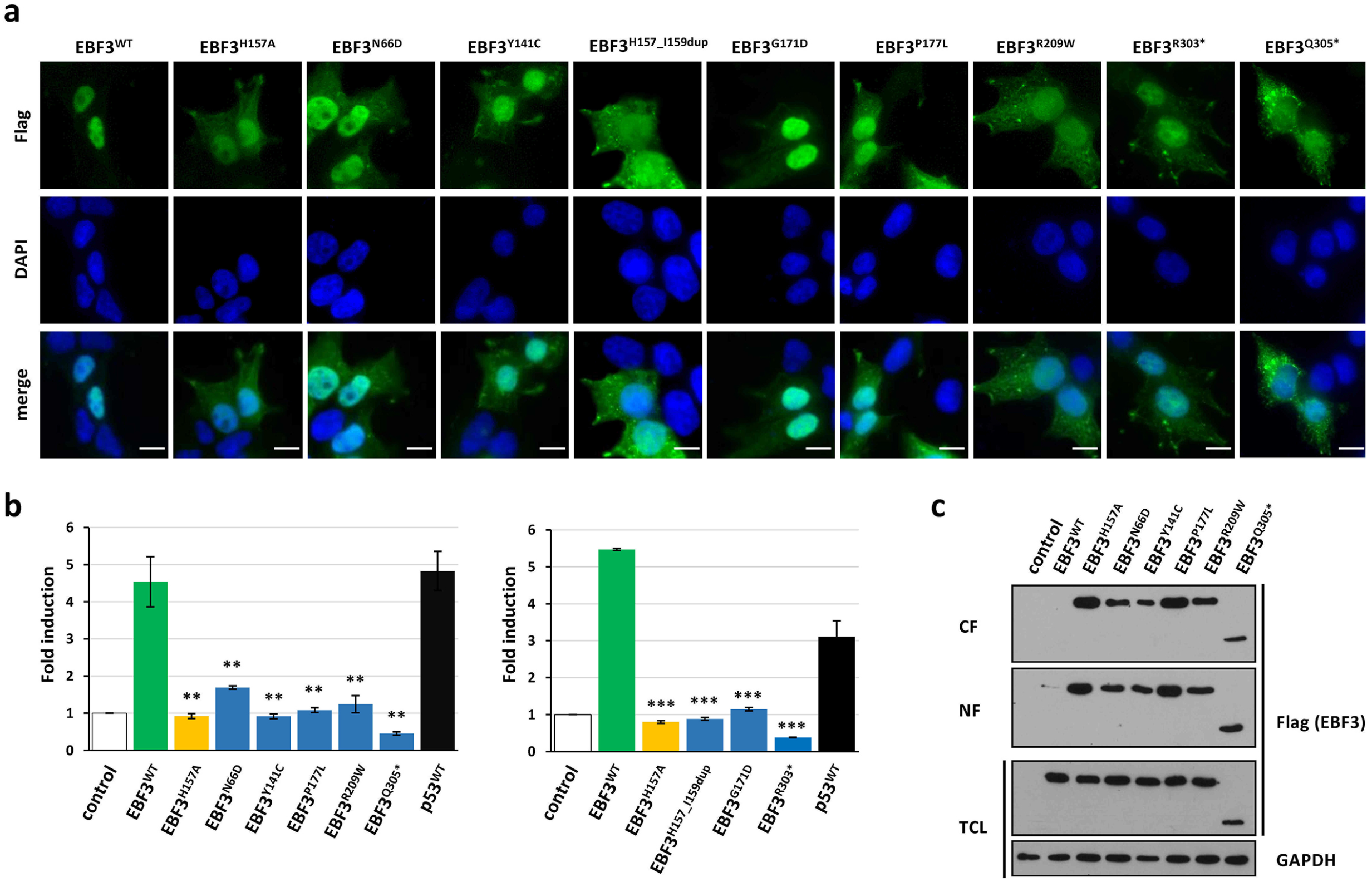
EBF3 mutants show impaired DNA-binding and altered subcellular localization. (**a**) Epifluorescence microscopy analysis was performed in HEK 293T cells transiently expressing wild-type or mutant EBF3 (green). Nuclei were stained with DAPI (blue). Wild-type EBF3 is exclusively localized in the nucleus, while the DNA-binding deficient mutant EBF3^H157A^ and the disease-associated mutants EBF3^N66D^, EBF3^Y141C^, EBF3^G171D^, EBF3^H157^_^I159dup^, EBF3^P177L^, EBF3^R209W^ and EBF3^R303*^ are also located in the cytoplasm. EBF3^Q305*^ is mainly localized in the cytoplasm. Representative images are shown. Bars correspond to 10 μm. (**b**) EBF3 mutants show impaired activation of luciferase reporter expression under the control of the *CDKN1A* (p21) promoter. HEK 293T cells were transiently transfected with EBF3 expression constructs or wild-type p53 as an internal control. Dual luciferase assays were done with the extracts of transfected cells 48 hours after transfection. Expression of wild-type EBF3 (green bar) and p53 (black bar) lead to a 4- to 5-fold elevated promoter activity compared with cells transiently transfected with empty vector (control; white bar). The DNA binding-deficient EBF3^H157A^ mutant (yellow bar) and all disease-associated EBF3 mutants (blue bars) showed a strongly reduced or no activation of the luciferase reporter. The normalized luciferase activity (mean ± s.d.) of three independent experiments is depicted as the fold induction relative to cells transfected with a control vector. All comparisons are in reference to wild-type EBF3, and *P* values were calculated using the two-sided Student’s *t* test. \*\**P* < 0.005. (**c**) EBF3 mutants are not tightly bound to chromatin. Transiently transfected HEK 293T cells were incubated with CSK buffer containing 0.1% Triton-X. The cytoplasmic extracts were removed and proteins precipitated. Cells were subsequently treated with CSK buffer supplemented with 0.5% Triton-X. Nuclear extracts were removed and proteins precipitated. Total cell lysate (TCL), cytoplasmic fraction (CF), and nuclear fraction (NF) were analyzed by SDS-PAGE and immunoblotting using an anti-Flag antibody. The mutant EBF3 proteins are present in both the CF and NF. In marked contrast, wild-type EBF3 is present in only minimal amount in the NF demonstrating exclusive nuclear localization and strong chromatin binding. Expression of EBF3 protein variants was monitored by immunoblotting using anti-flag antibody, and anti-GAPDH antibody was used to control for equal loading. Data shown are representative of four independent experiments.

To study interaction of EBF3 mutants with chromatin, we performed *in situ* subcellular fractionation. Transiently transfected HEK 293T cells, expressing either wild-type EBF3 or one of the mutants p.Arg209Trp, p.Asn66Asp, p.Pro177Leu, p.Tyr141Cys, and p.Gln305* were treated to extract cytoplasmic proteins followed by selective extraction of non-tightly chromatin-bound proteins and free protein aggregates within the nucleus. Detection of Flag-tagged EBF3 proteins by immunoblotting demonstrated all mutants to be present in both the cytoplasmic and the nuclear fraction (Fig. 2c). In contrast, wild-type EBF3 was absent in the fraction containing cytoplasmic proteins and only barely detectable in the nuclear fraction indicating that in contrast to EBF3 mutants the wildtype is tightly associated with chromatin (Fig. 2c).

To further test the hypothesis that the *EBF3* variants impact EBF3 DNA-binding and have regulatory consequences, we transfected SK-N-SH cells with *EBF3^WT^* and *EBF3^P177L^* cDNA overexpression constructs, stably selected the cells for integration and expression, and then performed RNA-sequencing. Overexpression of EBF3^WT^ or EB3F^P177L^ resulted in 1,712 or 509, respectively, differentially expressed transcripts relative to untransfected cells (FDR<0.05; Fig. 3a,b and **Supplementary Fig. 5**). Significantly enriched Gene Ontology terms associated with EBF3^WT^ expression include various neuronal development and signaling pathways, supporting a broad role for EBF3 in neurodevelopment (**Supplementary Fig. 6**). In contrast, EBF3^P177L^ did not yield as significant an enrichment in neurodevelopmental pathways. Overall, analyses of the SK-N-SH *EBF3^WT^* and *EBF3^P177L^* transcriptomes indicate that EBF3 targets a wide variety of genes and the p.Pro177Leu mutation leads to a reduction in the extent of EBF3-mediated gene regulation.

**Figure 3.**
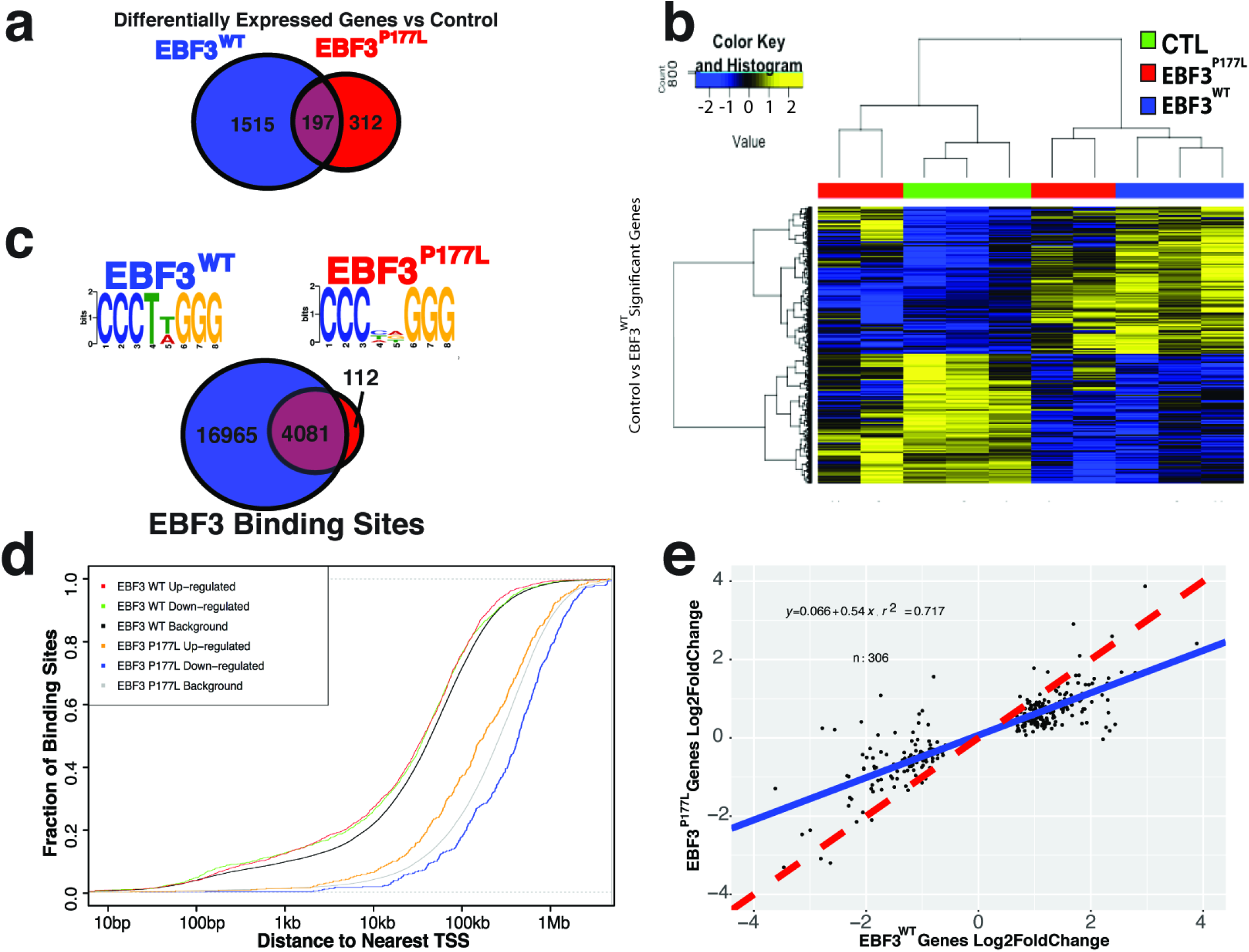
EBF3^P177L^ overexpression shows reduced transcriptome alteration and whole-genome EBF occupancy compared to EBF3^WT^. (**a**) EBF3^P177L^ reduces transcriptome alteration induced by *EBF3* expression relative to EBF3^WT^. Genes identified as significantly differentially expressed between SK-N-SH cells (control) and EBF3^WT^ and EBF3^P177L^ by DESeq2 with an adjusted p-value (FDR) < 0.05, with 197 shared differentially expressed genes between EBF3^WT^ and EBF3^P177L^. (**b**) A heatmap of DESeq2 variance-stabilized RNA-seq expression values comparing control (CTL), EBF3^WT^ and EBF3^P177L^ samples for genes determined to be significantly different between EBF3^WT^ and control SK-N-SH cells. (**c**) EBF3^P177L^ reduces genome-wide EBF3 binding-sites determined by ChIP-seq (bottom). ChIP-seq was performed using a high-affinity anti-FLAG antibody targeting the C-terminal 3X-FLAG epitope of the *EBF3* expression vectors. Most significant motifs with centrally enriched distribution for EBF3^WT^ and EBF3^P177L^ were identified using MEME-Suite (top). (**d**) EBF3^WT^ binding sites are enriched for closer proximity to nearest gene transcriptional start site (TSS) compared to EBF3^P177L^ binding sites. Cumulative Distribution Function (CDF) plot of EBF3^WT^ and EBF3^P177L^ shows binding sites and their distance to nearest GRCh37 gene TSS for genes identified as upregulated or downregulated (FDR<0.05, log2 fold change >0 and <0, respectively) compared to all genes. (e) EBF3^WT^ significantly differentially expressed genes with TSS within 5 kb of shared ChIP-seq binding sites exhibit relatively greater positive and negative log_2_ fold-change than the same genes in EBF3^P177L^ expression data. Linear regression of log_2_ fold-change values for these genes (n=306) exhibits a significantly downward-skewed slope of 0.54 (blue line) compared to the null expectation of perfect correspondence (slope = 1, dashed red line), indicating comparatively reduced alteration of expression for significant EBF3^WT^ genes by EBF3^P177L^.

To determine if EBF3^P177L^ affects binding of EBF3 across the genome, we performed chromatin immunoprecipitation coupled with massively parallel sequencing (ChIP-seq) in both *EBF3^WT^* and *EBF3^P177L^* transfected SK-N-SH cells. There were 21,046 binding sites identified for EBF3^WT^ and 4,193 binding sites for EBF3^P177L^, of which 4,081 were shared (Fig. 3c) – i.e., EBF3^P177L^ binds to a subset of EBF3^WT^ binding sites. MEME-Suite motif analysis^19^ identified an enrichment for the canonical EBF zinc binding motif in called peaks from both wild-type and mutant protein expressing lines (Fig. 3c). We then investigated whether or not EBF3^WT^ binding sites in our ChIP-seq data were enriched for closer proximity to gene transcriptional start sites (TSS) than EBF3^P177L^ binding sites. Compared to EBF3^P177L^, EBF3^WT^ binding sites were significantly more likely to reside near gene TSSs (K-S p<0.05) and genes differentially expressed in EBF3^WT^ were enriched for closer proximity to binding site peaks than non-significant genes (Fig. 3d). To determine if EBF3^WT^ proximal binding sites have some functional transcriptional consequence that is altered by the p.Pro177Leu mutation, we compared log2 fold-changes for EBF3^WT^ significantly differentially expressed genes with a TSS within 5 kb of shared EBF3^WT^ and EBF3^P177L^ binding sites. These genes had comparatively smaller log_2_ fold-changes in EBF3^P177L^ samples relative to EBF3^WT^ (Fig. 3e), indicating that the p.Pro177Leu mutation appears to lead to reduced transcriptional alteration. Taken together, these findings are consistent with EBF3 acting as a proximal regulator of transcription at cis-regulatory sequences and supports the hypothesis that EBF3^P177L^ has reduced function due to partial disruption of the DNA-binding domain.

In conclusion, our results show that *de novo* mutations disrupting the regulatory functions of the conserved neurodevelopmental transcription factor EBF3 underlie a new syndrome of intellectual disability, ataxia and facial dysmorphism. This phenotype, while rare, is a substantial contributor to intellectual disability. Two patients studied here were found in a single Clinical Sequencing Exploratory Research study^20^ on a series of 317 unrelated families. Additionally, 5 *de novo* protein altering variants (2 missense, 1 frameshift, 2 splice) in *EBF3* have recently been reported in a series of 4,293 families with individuals with developmental disorders, although neither these variants nor *EBF3* were discussed nor concluded to be pathogenic (preprint data available at http://biorxiv.org/content/early/2016/04/22/049056). Both missense variants are located in the DNA-binding domain of EBF3 and one, p.(Pro177Leu), is identical to a mutation in this study, indicating mutational recurrence. Combining these observations with our own analyses indicates that mutations in *EBF3* may underlie more than 1 in 1,000 individuals affected with otherwise unexplained developmental delay. Our study furthermore underscores the importance of data sharing and collaborative human genetics, with tools like GeneMatcher promoting the assembly of patient cohorts that enable identification of novel genetic contributions to disease and functionally annotate the human genome.

**URLs**. Online Mendelian Inheritance in Man (OMIM), http://www.omim.org/; NCBI Gene database, <u>http://www.ncbi.nlm.nih.gov/gene/</u>; ExAC Browser, http://exac.broadinstitute.org/; Exome Variant Server (EVS), http://evs.gs.washington.edu/EVS/; dbSNP, <u>http://www.ncbi.nlm.nih.gov/SNP/</u>; 1000 Genomes Project <u>http://www.1000genomes.org/</u>; Genic intolerance, <u>http://genic-intolerance.org/</u>; Genematcher, <u>http://www.genematcher.org/</u>; NCBI HomoloGene, <u>http://www.ncbi.nlm.nih.gov/homologene/</u>; Clustal Omega <u>http://www.ebi.ac.uk/Tools/msa/clustalo/</u>; SWISS-MODEL, <u>http://swissmodel.expasy.org/</u>; UCSF Chimera, <u>http://www.cgl.ucsf.edu/chimera/</u>; Protein Data Bank (PDB), <u>http://www.rcsb.org/pdb/home/home.do</u>; database for nonsynonymous SNPs’ functional predictions (dbNSFP), <u>https://sites.google.com/site/jpopgen/dbNSFP</u>;

## ACKNOWLEDGEMENTS

We are grateful to the patients and their families who contributed to this study. We would like to thank members of the HudsonAlpha CSER team, especially Shirley Simmons, Kelly East, Whitley Kelley, Candice Finnila, David Gray, Michelle Amaral, and Michelle Thompson, Ryne Ramaker for help with sequencing data analysis, Verena Kolbe for skillful technical assistance, Hans-Jürgen Kreienkamp for help with structural analysis and critical reading, Stefan Kindler and Claudia Schob for helpful suggestions and technical advice with the luciferase assay, and the UKE Microscopy Imaging Facility (UMIF) for technical support. This work was supported by grants from the US National Human Genome Research Institute (NHGRI) (UM1HG007301), the US National Institutes of Health (1R21NS094047-01), the NHGRI/National Heart Lung and Blood Institute (NHLBI) [No. HG006542 to the Baylor-Hopkins Center for Mendelian Genomics (BHCMG)], the UAB MSTP (NIH-NIGMS 5T32GM008361-21 to A.H.), the Simons Foundation (to W.K.C.), and the Deutsche Forschungsgemeinschaft (KO 4576/1-1 and 1-2 to F.K. and KU 1240/10-1 to K.K.). W.-L.C. was supported by The Cancer Prevention & Research Institute of Texas (CPRIT) training Program RP140102.

## AUTHOR CONTRIBUTIONS

F.L.H. performed WES, exome data interpretation, mutation validation and segregation analysis for family 1, cloning, structural analysis, immunocytochemistry, microscopy, transactivation assay, *in situ* subcellular fractionation and wrote the manuscript. K.M.G., A.S. and A.D. recruited and clinically characterized subjects of family 1, collected biological samples, evaluated and summarized clinical data and wrote the manuscript. A.A.H. performed the ChIP-seq, RNA-seq, and related experiments and analyses and wrote the manuscript. F.K. performed WES and data interpretation for family 1. M.A. performed bioinformatics WES data processing and analysis for family 1. L.B. and M.T. recruited and clinically characterized subjects of family 2 and collected biological samples. L.M.B. and S.C. recruited and clinically characterized subjects of family 3 and collected biological samples. E.J.L and M.B. recruited and clinically characterized subjects of families 4 and 5 and collected biological samples. K.M.B. and S.M.H. performed the genomic analyses to identify the *EBF3* variants in families 4 and 5. W.K.C. recruited and clinically characterized subjects of family 6 and collected biological samples. M.K.E., Z.C.A., W.-L.C. and M.B. performed rare variant analysis of WES data in families 7 and 8 and segregation studies. M.D.-B. recruited and clinically characterized subjects of family 7 and collected biological samples. A.W.E.-H. and M.A.M.S. recruited and clinically characterized subjects of family 8 and collected biological samples. S.B. contributed to phenotype and biological samples collection, and to WES as investigator of the study for family 9. B.C. performed bioinformatic analyses and evaluated and interpreted WES data for family 9. B.I. recruited and clinically characterized subjects of family 9 and collected biological samples. S.K. contributed to collection of family 9 information, evaluated and interpreted WES data and wrote the manuscript. J.R.L. was in charge of the experimental design and analyses of data and supervised studies for families 7 and 8. G.M.C. and R.M.M. designed the study on families 4 and 5 (WES, ChIP-seq, RNA-seq, and related experiments), guided implementation across all components and wrote the manuscript. K.K. conceived the project, analyzed and interpreted the data, and wrote the manuscript. All authors contributed to and approve of the manuscript.

## Material and Methods

### Subjects

Ten subjects with intellectual disability and additional clinical manifestations were included in the study. Clinical features are summarized in **Supplementary Table 4**. Informed consent for DNA storage and genetic analyses was obtained from the parents of all subjects, and genetic studies were approved by all institutional review boards of the participating institutions. Permission to publish photographs was given for all subjects shown in Figure 1c.

#### Exome sequencing and sequence data analysis

Targeted enrichment and massively parallel sequencing were performed on genomic DNA extracted from circulating leukocytes. Enrichment of the whole exome was performed according to the manufacturer’s protocols using the Nextera Enrichment Kit (62 Mb) (Illumina) for subjects 1 and 2 and their brother (sibling 1)^1^, and the SureSelect Clinical Research Exome kit (54 Mb; Agilent) for subject 10. Captured libraries were then loaded onto the HiSeq2000 or 2500 platform (Illumina). Trimmomatic was employed to remove adapters, low quality (phred quality score < 5) bases from the 3’ ends of sequence reads^2^. Cutadapt was used for subject 10. Reads shorter than 36 bp were subsequently removed. Further processing was performed following the Genome Analysis Toolkit’s (GATK) best practice recommendations. Briefly, trimmed reads were aligned to the human reference genome (UCSC GRCh37/hg19) using the Burrows-Wheeler Aligner (BWA mem v0.7.12). Duplicate reads were marked with Picard tools (v1.141). GATK (v3.4) was employed for indel realignment, base quality score recalibration, calling variants using the HaplotypeCaller, joint genotyping, and variant quality score recalibration. AnnoVar (v2015–03–22) was used to functionally annotate and filter alterations against public databases (dbSNP138, 1000 Genomes Project, and ExAC Browser). Exonic variants and intronic alterations at exon-intron boundaries ranging from −10 to +10, which were clinically associated and unknown in public databases, were retained. Whole-exome sequencing and data analysis for families 2, 3, and 6 was performed at GeneDx as described previously^3^. For families 7 and 8, WES and data analysis were performed in Human Genome Sequencing Center (HGSC) at Baylor College of Medicine, according to the previously described protocol and using notably the VCRome capture reagent^4–6^; WES targeted the coding exons of ~20,000 genes with 130X average depth-of-coverage and greater than 95% of the targeted bases with >20 reads^4–6^.

Subjects 5 and 6 (families 4 and 5) were identified as part of a trio-based clinical sequencing study at HudsonAlpha, which to date has performed sequencing on 349 affected probands from 317 families. Genomic DNA was isolated from peripheral blood leukocytes and exome sequencing was conducted to a median depth of ~65X with at least 80% of targeted bases covered at 20X. Exome capture was completed using Nimblegen SeqCap EZ Exome version 3 and sequencing was conducted on the Illumina HiSeq 2000. Reads were aligned to reference hg19 using bwa (0.6.2)^7^. GATK best practice methods^8^ were used to identify variants, with samples called jointly in batches of 10–20 trios. Maternity and paternity of each parent was confirmed by whole exome kinship coefficient estimation with KING^9^. *De novo* variants were identified as heterozygous variant calls in the proband in which there were at least 10 reads in both parents and the proband, an alternate allele depth of at least 20% in the proband and less than 5% in each parent, and a minor allele frequency <1% in 1000 Genomes and ExAC. All candidate *de novo* variants that either affected a protein or had a scaled CADD score > 10 (ref. ^10^) were manually reviewed, as were variants similarly identified using X-linked, compound heterozygote, and recessive inheritance filters. In each patient described, no other variants were identified as potentially disease-causal.

#### Variant validation

Sequence validation and segregation analysis for all candidate variants were performed by Sanger sequencing. Primer pairs designed to amplify selected coding exons of the *EBF3* gene and exon-intron boundaries (NM_001005463.2) and PCR conditions are available on request. For families 1 and 9, amplicons were directly sequenced using the ABI BigDye Terminator Sequencing Kit (Applied Biosystems) and an automated capillary sequencer (ABI 3500; Applied Biosystems). Sequence electropherograms were analysed using Sequence Pilot software (JSI medical systems). For families 4, 5, 7 and 8, Sanger confirmation of variants, including confirmation of absence from both biological parents, was performed by a CAP/CLIA-certified diagnostic laboratory. Genotyping was carried out with the AmpFl STR SGM plus Kit (Applied Biosystems) to confirm paternity and maternity. The effects of the variants were assessed with Combined Annotation-Dependent Depletion (CADD)^11^ and by determining the GERP++ scores^12, 13^.

#### RNA isolation, cDNA synthesis, cloning and colony PCR

Total RNA was extracted (RNeasy Mini kit, Qiagen) from cultured primary fibroblasts obtained from a skin biopsy of subjects 1 and 2. 1 µg total RNA was reverse transcribed (Superscript™ III RT, ThermoFisher) using random hexamers as primers, and 1 μl of the reverse transcription reaction was utilized to amplify a 320-bp *EBF3* cDNA fragment encompassing the c.625C>T variant (forward primer 5′-ACCCACGAGATCATGTGCAG-3′, reverse primer 5′-CTGGGACTGATGGCCTTG-3′). The PCR product was directly sequenced.

Exon 7 of *EBF3* and adjacent intronic sequences were amplified from leukocyte- and buccal cell-derived DNA of the mother of subjects 1 and 2. The PCR product was cloned into the pCR2.1 TOPO TA Cloning Vector (ThermoFisher). Individual *E.coli* clones were subjected to colony PCR followed by Sanger sequencing to haplotype determination.

#### Structural analysis

The three-dimensional structure of the DNA-binding domain of wildtype and EBF3 mutants (amino acids 50–251) was obtained by means of homology modeling using the SWISS-MODEL web-based service^14^. The crystallographic structure at 2.8–Å resolution of EBF1 bound to DNA (PDB entry 3MLP) was used as a template^15^. The selected template ensured the best homology score (95.9%). The coordinates of the DNA backbone were taken from the crystal structure of the double-stranded DNA bound to EBF1^15^. Molecular graphics were developed with UCSF Chimera software^16^.

#### Plasmid construction and mutagenesis

The coding region of human wild-type *EBF3* (NM_001005463.2) was amplified by using *EBF3*-specific PCR primers and cDNA of human fetal brain as template. The forward primers were designed with a 5′-CACC overhang and lacked the start codon sequence. Purified PCR products were cloned into pENTR/D-TOPO vector (ThermoFisher) according to the manufacturer’s protocol. *EBF3* single mutations encoding p.His157Ala, p.Asn66Asp, p.Tyr141Cys, p.His157_Ile159dup, p.Gly171Asp, p.Pro177Leu und p.Arg209Trp were introduced into *EBF3* cDNA with the QuikChange II Site-Directed Mutagenesis Kit (Agilent Technologies). Subsequently, N-terminally FLAG-tagged EBF3 constructs were generated by transferring the coding region into the destination vector pFLAG-CMV4-cassetteA^17^. The coding region of the mutants EBF3^Q305*^ and EBF3^R303*^ and of p53^WT^ was amplified by using specific PCR primers and pFLAG-CMV4-EBF3^WT^ and LeGO-iG2-Puro+−p53 as templates, respectively. The LeGO-iG2-Puro+−p53 construct was kindly provided by Kristoffer Rieken (University Medical Center Hamburg-Eppendorf, Hamburg, Germany). Cloning into pENTR/D-TOPO and destination vectors was performed as described above. For RNA-sequencing and ChIP-seq experiments EBF3^WT^ and EBF3^P177L^ cDNA constructs were designed and ordered as synthetic dsDNA gBlocks (IDT) and cloned into a pCMV6–3xFlag-2A-Neomycin vector (modified from pCMV6-AC-GFP, Origene) using Gibson Assembly (NEB) to yield the desired EBF3–3xFlag expression vectors (**Supplementary Fig. 7**). Full vector and cDNA sequences are available on request. All constructs were sequenced for integrity.

#### Cell culture and transfection

HEK 293T (human embryonic kidney) cells and primary fibroblasts were cultured in Dulbecco’s modified Eagle medium (DMEM; ThermoFisher) supplemented with 10% fetal bovine serum (FBS; GE Healthcare) and penicillin-streptomycin (100 U/mL and 100 mg/mL, respectively; ThermoFisher). HEK 293T cells were transfected using TurboFect (ThermoFisher) with a DNA (μg): TurboFect (μl) ratio of 1:2 for immunocytochemistry and *in situ* fractionation experiments and of 1:1 for transactivation assays.

SK-N-SH (ATCC HTB-11) cells (which do not endogenously express EBF3), were obtained from ATCC and grown under recommended growth conditions. EBF3 constructs were transfected into SK-N-SH cell using Nucleofection (Lonza) kit V using manufacturer’s instructions, with the following modifications: 5 ug of *EBF3^WT^* or *EBF3^P177L^* vector was transfected per 1×10^6^ cells, and three transfections were pooled into one biological replicate for a total of four biological replicates per vector. Cells were selected with 400 ug/mL G418 (Invitrogen, 10131035) for two weeks and then with 200 ug/mL for a further 2–3 weeks to generate SK-N-SH cells with stable EBF3-3xFlag expression. Biological replicates were maintained under selection as polyclonal pools and expanded for use in functional genomic experimentation.

#### Immunocytochemistry

HEK 293T cells were cultivated on glass coverslips coated with Poly-L-lysine (Sigma Aldrich) and transiently transfected with *EBF3* expression constructs. 24 h after transfection, cells were fixed with 4% paraformaldehyde (Sigma-Aldrich) in PBS. After treatment with permeabilization/blocking solution (2% BSA; 3% goat serum; 0.5% Nonidet P40 in PBS), cells were incubated in antibody solution (3% goat serum; 0.1% Nonidet P40 in PBS) containing mouse monoclonal anti-FLAG M2 antibody (1:200 dilution; clone: F-3165; Sigma-Aldrich). Cells were washed with PBS and incubated with Alexa Fluor-488 coupled goat anti-mouse IgG (1:1000 dilution; ThermoFisher). After extensive washing with PBS, cells were embedded in mounting solution (ProLong Diamond Antifade Mountant with DAPI; ThermoFisher). Cells were analyzed with Olympus IX-81 epifluorescence microscope equipped with a 60x Plan Apo N oil immersion objective lens.

#### Transactivation assay

For transactivation assays, HEK 293T cells were transiently transfected to express the construct(s) of interest, together with the pGL2-p21 (CDKN1A) promoter-Luc^18^ and pRen using a 1:1:3 ratio of pREN(2 µg):pGL2-p21 promoter-Luc(2 µg):pFLAG-CMV4-EBF3 or pFLAG-CMV4-p53^WT^(6 µg) expression constructs. For coexpression experiments with EBF3^WT^ and mutant EBF3, a 1:1:2:2 ratio of pREN:pGL2-p21 promoter-Luc:pFLAG-CMV4-EBF3^WT^:pFLAG-CMV4-EBF3^mut^ was used. For titration experiments (with wild-type EBF3 and either EBF3^N66D^ or EBF3^Q305*^) cells were transfected with a constant amount of wild-type EBF3 (4 µg) and an increasing amount of the EBF3 mutant constructs. Transactivation assays performed to analyze the dose-dependent activation of wild-type EBF3 on the pGL2-p21 promoter-Luc construct documented that a maximum activation was attained with 6 to 8 µg of wild-type EBF3 construct (Supplementary Fig. 8). The pGL2-p21 promoter-Luc was a gift from Martin Walsh (Addgene plasmid #33021) and encodes *Photinus* luciferase. The eukaryotic expression vector pREN is a derivate of the pFiRe-basic^19^ containing a recombinant gene that is under the control of cytomegalovirus immediate early promoter and encodes *Renilla* luciferase kindly provided by Stefan Kindler (University Medical Center Hamburg-Eppendorf, Hamburg, Germany). The Dual-Luciferase Reporter Assay System (Promega) was performed according to the manufacturer’s protocol with cell extracts prepared 48 h after transfection. Data were normalized to the activity of *Renilla* luciferase, and basal promotor activity for transfection with pFLAG-CMV4-cassetteA (control vector) was considered 1. All assays were performed in triplicate.

#### *In situ* subcellular fractionation

3.5 × 10^5^ HEK 293T cells were seeded on 6-well plates and incubated under normal growth condition overnight. 24 h after transfection of cells with EBF3 expression constructs or pFLAG-CMV4-cassetteA (control vector), *in situ* subcellular fractionation was performed as described previously^20^. Briefly, cells were washed twice with ice-cold PBS, followed by a 1 min incubation on ice with 500 µL CSK buffer (10 mM PIPES, pH 6.8; 100 mM NaCl; 300 mM sucrose; 3 mM MgCl_2_; 1 mM EGTA) supplemented with 0.1% Triton-X. The cytoplasmic fraction (CF) was removed and proteins were precipitated with 1 M (NH_4_)_2_SO_4_ at 4°C under rotating conditions. Subsequently, cells were washed with ice-cold PBS and then incubated for 20 min on ice with 500 µl CSK buffer supplemented with 0. 5% Triton-X. The nuclear fraction (NF) encompassing nuclear proteins and proteins loosely bound to chromatin was removed and proteins were precipitated. Total cell lysate (TCL) was prepared from a separate well by adding 750 µl TES buffer (1% SDS; 2 mM EDTA; 20 mM Tris-HCl, pH 7.4) for 1 min at room temperature. TCL was removed and proteins were precipitated. Samples were centrifuged at 14,000 rpm for 30 min at 4°C and protein pellets were resuspended in 1x SDS loading buffer. Proteins were separated on 12.5% SDS-polyacrylamide gels and transferred to PVDF (polyvinylidene difluoride) membranes. Following blocking (20 mM Tris-HCl, pH 7.4; 150 mM NaCl; 0.1% Tween-20; 5% nonfat dry milk) and washing (20 mM Tris-HCl, pH 7.4; 150 mM NaCl; 0.1% Tween-20) membranes were incubated in primary antibody solution (20 mM Tris-HCl, pH 7.4; 150 mM NaCl; 0.1% Tween-20) containing mouse monoclonal anti-FLAG M2 peroxidase conjugate (1:50,000 dilution; Sigma-Aldrich). After final washing, immunoreactive proteins were visualized using the Immobilon Western Chemiluminiescent HRP substrate (Sigma-Aldrich). For control of equal loading, TCL was analyzed using mouse anti-GAPDH antibody (1:10,000 dilution; Abcam) in primary antibody solution containing 0.5% nonfat dry milk. Membranes were washed and incubated with peroxidase-coupled anti-mouse IgG (1:10,000 dilution; GE Healthcare).

#### RNA-seq

Total RNA was harvested from control SK-N-SH cells and SK-N-SH cells stably expressing *EBF3^WT^* or *EBF3^177L^* using the Norgen Total RNA Preparation Kit (Norgen Biotek). cDNA was prepared using the ThemoFisher High-Capacity RNA-to-cDNA Kit (ThermoFisher). RNA-seq libraries were prepared with Nextera DNA Library Sample Prep Kit using established protocols^21^. Libraries were sequenced on an Illumina HiSeq2500 with 50-bp paired-end sequencing. All statistical analyses were performed in R (Version 3.2.1). RNA-seq reads were processed using a custom pipeline implementing aRNApipe^22^ to perform low-quality read filtering, adaptor trimming, alignment with STAR^23^, and generate the final count table for samples passing an unambiguously-mapped alignment rate cutoff of at least 40%. To perform differential gene expression analysis, we used the R DESeq2 package^24^ and likelihood ratio test (LRT) hypothesis testing with an adjusted p-value (FDR) cutoff of 0.05. Heatmap and gene expression boxplots were generated using variance stabilized counts generated with DESeq2. GO Term enrichment was performed using the online tool g:Profiler^25^ with all GO term annotation categories.

#### ChIP-seq

Currently, fewer than 10% of antibodies tested for use in ChIP-seq have met quality control metrics set by ENCODE^26, 27^, and lack of high-affinity antibodies for proteins bound to fixed chromatin has been a major impediment to investigation of neurodevelopmental TF binding^28^. Additionally, available anti-EBF3 antibodies have been found to have some degree of cross reactivity with other EBF family members, potentially limiting interpretability of specific family member binding sites^29^. Although *EBF1* and *EBF2* are not expressed in the SK-N-SH cell line, *EBF4* is expressed highly. Thus, we chose to perform ChIP against tagged EBF3 protein rather than rely on native anti-EBF3 antibodies (high-affinity FLAG antibody that targets the C-terminal 3X-FLAG epitope of the *EBF3* cDNA constructs). *EBF3^WT^* and *EBF3^P177L^* genome-wide binding site identification was performed with ChIP-seq using established methods^30^. Briefly, 20 × 10^6^ cells for two biological replicates for each vector were crosslinked in 1% formaldehyde, sonicated to fragment chromatin using the BioRuptor Twin Sonicator (Diagenode), and immunoprecipitated using monoclonal anti-FLAG M2 antibody (Sigma). Samples were reverse-crosslinked and recovered DNA was used as input for Illumina sequencing library preparation. Libraries were sequenced on an Illumina NextSeq with single-end 150 bp sequencing. ChIP-seq reads were aligned using the BWA aligner to hg19 and peaks were identified for each replicate using MACS2.1.0^31^ with an –mfold cutoff of (10,30). Replicate overlapping peaks were merged using BEDTools^32^ to generate final peak lists used in downstream analyses. Motif identification was performed using MEME-Suite^33^ with HOCOMO (v10) with final peaks sequences trimmed to a centered 100 base pair window.

#### Statistical analysis

Differences in the distribution of continuous variables between groups were evaluated for statistical significance using two-paired Student’s *t* test. In all comparisons, *P* values of ≤0.05 were considered to be statistically significant.

